# Neural correlates of blood flow measured by ultrasound

**DOI:** 10.1101/2021.03.31.437915

**Authors:** Anwar O. Nunez-Elizalde, Michael Krumin, Charu Bai Reddy, Gabriel Montaldo, Alan Urban, Kenneth D. Harris, Matteo Carandini

## Abstract

Functional ultrasound imaging (fUSI) is a popular method for measuring blood flow and thus infer brain activity, but it relies on the physiology of neurovascular coupling and requires extensive signal processing. To establish to what degree its trial-by-trial signals reflect neural activity, we performed simultaneous fUSI and neural recordings with Neuropixels probes in awake mice. fUSI signals strongly correlated with the slow (<0.3 Hz) fluctuations in local firing rate, and were closely predicted by the smoothed firing rate of local neurons, particularly putative inhibitory neurons. The optimal smoothing filter had width ~3 s, matched the hemodynamic response function of awake mouse, was invariant across mice and stimulus conditions, and similar in cortex and hippocampus. fUSI signals also matched neural firing spatially: firing rates were as highly correlated across hemispheres as fUSI signals. Thus, hemodynamic signals measured by ultrasound bear a simple and accurate relationship to neuronal firing.

## Introduction

Functional ultrasound imaging (fUSI) is an increasingly popular method for studying brain function (Deffieux et al., 2018; Edelman and Macé, 2021; Macé et al., 2011; Rabut et al., 2020). fUSI is appealing because it estimates changes in cerebral blood volume with high resolution, resolving spatial features in the order of ~100 μm up to a depth of ~2 cm (Macé *et al*., 2011). It is thus used to study how the activity of brain regions depends on sensory stimuli, internal state, and behavior, in multiple species including mice (Aydin et al., 2020; Boido et al., 2019; Brunner et al., 2020; Ferrier et al., 2020; Koekkoek et al., 2018; Macé et al., 2018; Sans-Dublanc et al., 2021), rats (Bergel et al., 2018; Bergel et al., 2020; Gesnik et al., 2017; Macé *et al*., 2011; Osmanski et al., 2014; Provansal et al., 2021; Rahal et al., 2020; Sieu et al., 2015; Urban et al., 2015), marmosets (Zhang et al., 2021), ferrets (Bimbard et al., 2018), and macaques (Blaize et al., 2020; Dizeux et al., 2019). In a small animal like a mouse, fUSI can image the whole brain, yielding measurements that may parallel those obtained in humans with functional magnetic resonance imaging (fMRI).

However, the relationship between fUSI signals and neural activity is indirect, as it relies on multiple intermediate steps: the physiology of neurovascular coupling, the physics of ultrasound sensing, and the mathematics of the subsequent signal processing. Neurovascular coupling links neuronal firing to changes in blood oxygenation, flow, and volume through processes that have been extensively studied (Attwell and Iadecola, 2002; Drew, 2019; Hamel, 2006; Hillman, 2014; Iadecola and Nedergaard, 2007; Nair, 2005; Pisauro et al., 2013; Turner et al., 2020; Winder et al., 2017). Movement of blood, in turn, causes a frequency shift in ultrasound echoes that can be measured through power Doppler ultrasound sensing (Rubin et al., 1995; Rubin et al., 1994). Finally, and crucially, the power Doppler signals must be distinguished from multiple, large sources of noise -- such as tissue movement -- through multiple steps of signal processing. These steps typically include temporal binning, power estimation, temporal high-pass filtering and spatiotemporal clutter filtering by removing the largest principal components (Baranger et al., 2018; Demené et al., 2015; Macé *et al*., 2011; Macé et al., 2013). Small changes in this procedure can profoundly affect the inferred neural signals (e.g., Demené *et al*., 2015). Yet this procedure has not been verified with simultaneous recordings of neuronal firing rates in the awake brain. Indeed, it is unclear to what degree, and at what temporal and spatial scales, fUSI signals truly measure neural firing on a trial-by-trial basis.

At first sight, fUSI signals may appear noisy, with large fluctuations over short time scales (e.g. >10% over a few seconds) that vary across trials (e.g., Brunner *et al*., 2020), and it is not clear to what extent these fluctuations are due to the process of measurement and analysis of fUSI signals, or to the underlying neural activity. Indeed, neural activity exhibits endogenous, ongoing fluctuations that are strongly correlated across neurons (Schölvinck et al., 2015), associated with changes in brain state and body movement (Drew et al., 2019; Musall et al., 2019; Stringer et al., 2019), and are highly correlated across hemispheres (Drew *et al*., 2019; Fox et al., 2007; Fox et al., 2006; Mohajerani et al., 2010; Shimaoka et al., 2019). Perhaps the apparently noisy fUSI signals reflect these structured fluctuations in neural activity. Indeed, fUSI signals approximately resemble simultaneously recorded local field potentials (Aydin *et al*., 2020; Bergel *et al*., 2018; Bergel *et al*., 2020; Sieu *et al*., 2015), which in turn reflect local neuronal firing (Buzsáki et al., 2012; Katzner et al., 2009).

Moreover, it is not clear whether the neural component of fUSI signals reflect neuronal spiking through a simple linear relationship and if this relationship differs across brains and brain regions. To a first approximation, neurovascular coupling is a linear process: hemodynamic signals can be predicted from neuronal firing by smoothing firing rates with a hemodynamic response function (Boynton et al., 1996; Cardoso et al., 2019; Devor et al., 2005; Drew, 2019; Heeger and Ress, 2002; Lima et al., 2014; Logothetis et al., 2001; Martindale et al., 2003; Pisauro *et al*., 2013). The next step might also be linear: fUSI signals can be predicted from (separately measured) hemodynamic signals (red blood cell velocity) through another transfer function (Aydin *et al*., 2020; Boido *et al*., 2019). Because a series of linear operations is itself linear, the relationship between fUSI signals and neuronal firing may result from a simple convolution with a linear filter. Furthermore, this relationship may be fixed across brain regions and types of activity.

Here we answer these questions with simultaneous measurements of spikes and fUSI signals in awake mice. We performed these experiments in the awake brain to avoid the detrimental effects of anesthesia on neurovascular coupling (Pisauro *et al*., 2013) and on the function of inhibitory circuits (Haider et al., 2013). The results indicate that fUSI signals are closely related to neuronal firing, and especially the firing of putative inhibitory neurons, and that the relationship between the two is well summarized by convolution with a hemodynamic response function. The transfer function acts as a low-pass filter, so the relationship between fUSI signals and neuronal firing becomes progressively more accurate at slower time scales. Neural activity explains why fUSI signals correlate strongly across space and even across hemispheres: these correlations reflect true shared fluctuations in neural activity across brain locations and hemispheres.

## Results

To measure neuronal firing during fUSI, we recorded with Neuropixels probes during sensory and spontaneous activity. For each mouse, we determined the location of primary visual cortex (V1) by aligning fUSI images to the Allen Institute Brain Atlas (Wang et al., 2020) using a vascular atlas as an intermediate reference (Todorov et al., 2020). In each session, we inserted a Neuropixels probe (Jun et al., 2017) in a parasagittal trajectory and acquired a fUSI image coronally (**Figure 1**A,B).

**Figure 1.**
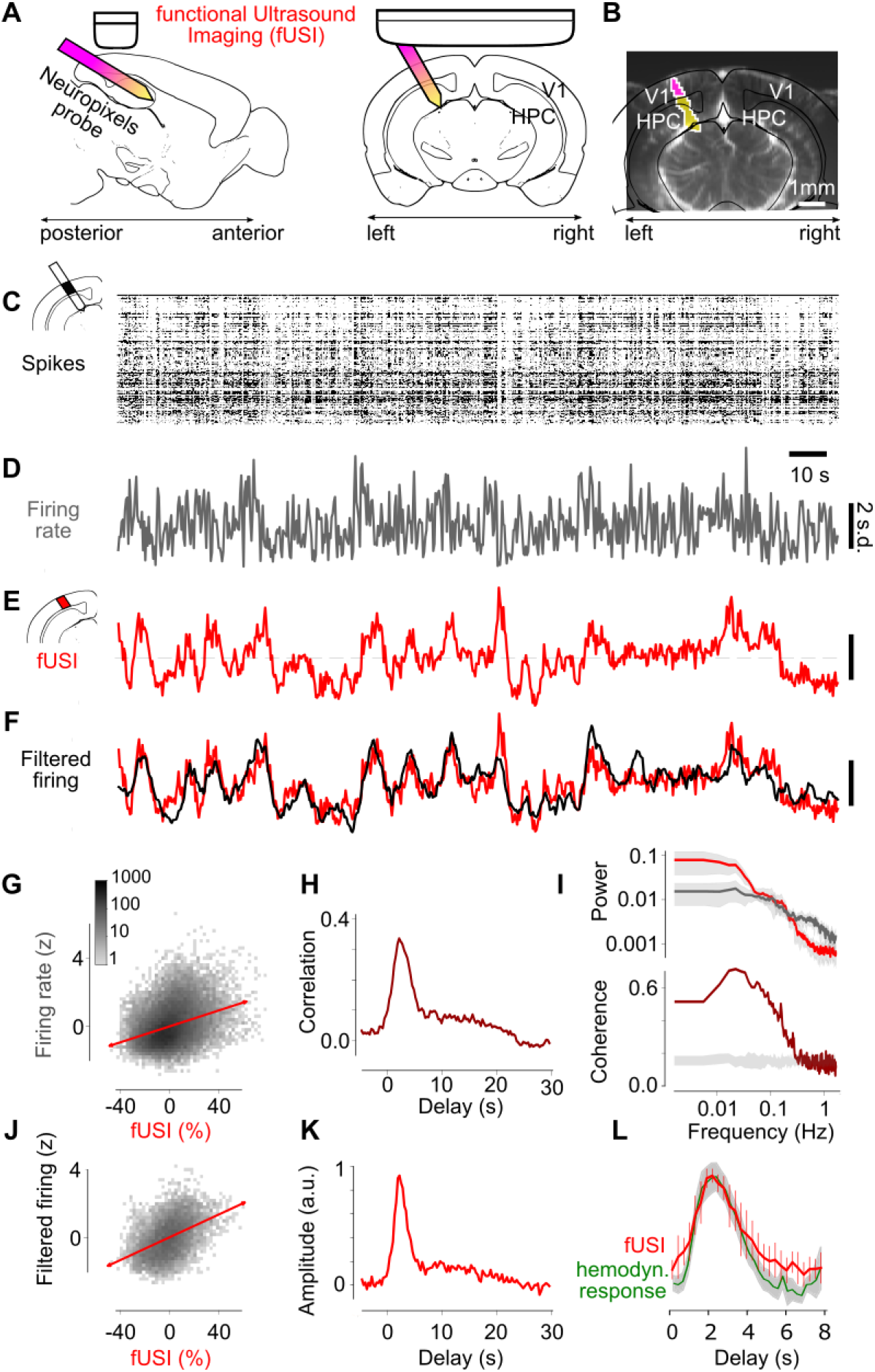
fUSI signal reflects temporally filtered firing rate during spontaneous activity. (**A**) Schematic of simultaneous fUSI and electrophysiological recordings, showing primary visual cortex (V1) and hippocampus (HPC). (**B**) Coronal fUSI slice with the location of the Neuropixels probe passing through this plane (*purple*) and in front of it (*yellow*). (**C**) Spikes recorded in V1 in an example recording, as a function of time and recording depth. (**D**) The resulting mean firing rate. (**E**) fUSI signal measured simultaneously in the same location (average over 51 voxels). (**F**) Smoothing the firing rate with the optimal filter (shown in K) yields good predictions (*black*) of the fUSI signals (*red*). (**G**) Comparison of fUSI signals and firing rate measured 2.1 s earlier (the optimal value), with best fitting line indicating correlation (red). 34 recordings in 5 mice. (**H**) Cross-correlation between firing rate and fUSI signal, averaged across 34 recordings in 5 mice. (**I**) Power spectra (top) and spectral coherence (bottom) of firing rate and fUSI, averaged across recordings. The gray bands in the top plot show 1 median absolute deviation (m.a.d.). The gray band in the bottom plot shows coherence of randomly circularly shifted traces. (**J**) Comparison of fUSI signals and filtered firing rate. (**K**) Optimal linear filter across recordings, obtained with cross-validation. Median of 34 recordings in 5 mice. (**L**) The filter (red) resembles the hemodynamic response function measured in awake mice (*green*, from (Pisauro *et al*., 2013)). Error bars show ± m.a.d. of 34 recordings in 5 mice.

Mice were awake and generally alert, as confirmed by measures of pupil dilation and whisker movements (McGinley et al., 2015; Reimer et al., 2014) (**Suppl. Figure 1**). They viewed a gray screen (to measure spontaneous activity) or flashing checkerboards (to measure visual responses). All recordings were repeated after moving the fUSI transducer ~0.4 mm to an adjacent coronal slice (35 slices per session). At the end of a session, we determined the location of the probe in the fUSI images by slowly extracting it while detecting its movement with fUSI (**Figure 1**B). To process fUSI signals we used established procedures (Demené *et al*., 2015; Macé *et al*., 2011), so that our results could be directly compared to the literature.

The fUSI signals from visual cortex during spontaneous activity resembled a delayed and smoothed version of the firing rate measured in the same location. After spike sorting, we computed the mean firing rate in all neurons (both single- and multi-unit clusters) recorded at the sites that intersected the fUSI slice (**Figure 1**C,D). This firing rate resembled the fUSI signal measured in the corresponding voxels (**Figure 1**E). The correlation between firing rate (delayed by 2.1 s) and fUSI signals was ρ = 0.34 ± 0.08 (median ± median absolute deviation, m.a.d., 34 recordings in 5 mice, **Figure 1**G). The delay between the two signals was estimated by plotting the cross-correlation as a function of time, which peaked at a delay of 2.1 ± 0.3 s, with full-width at half-height of 3.6 ± 0.6 s (± m.a.d., 34 recordings in 5 mice, **Figure 1**H).

Firing rate and fUSI signals were strongly correlated at low frequencies and became progressively less correlated at higher frequencies. To estimate the correlation between fUSI and firing rate as a function of frequency, we computed their spectral coherence, i.e., their correlation as a function of frequency (**Figure 1**I). Coherence was highest between 0.01 and 0.1 Hz, with a median correlation of 0.59 ± 0.03 (m.a.d., 34 recordings in 5 mice), and gradually fell to chance levels (coherence of 0.14 ± 0.03) at a frequency of 0.32 Hz. These results indicate that low frequency fluctuations in fUSI are mostly neural in origin, whereas fluctuations at higher frequencies are unrelated to neural activity and might thus best be discarded.

The precise relationship between fUSI signals and firing rate was well described by convolution with a linear filter. The cross-correlation between two signals reflects not only their interaction but also their individual autocorrelations, which are substantial in both firing rates and fUSI signals. To obviate this problem, we estimated the optimal filter that relates the two through convolution (Boynton *et al*., 1996; Pisauro *et al*., 2013), using cross-validated ridge regression (Hoerl and Kennard, 1970). Smoothing the firing rate with this filter yielded a prediction that closely matched the fUSI signal (**Figure 1**F). The filtered firing rate and the fUSI signal were highly correlated: in held-out data, the median correlation between the two was ρ = 0.49 ± 0.13 (m.a.d., 34 recordings in 5 mice, **Figure 1**J).

The filter relating fUSI signals to firing rate resembled the hemodynamic response function characteristic of awake mouse cortex. As expected, the estimated filter peaked with the same delay as the cross-correlations (2.1 ± 0.3 s, median ± m.a.d.), but it had a faster time-course. Its full width at half-height was 2.9 ± 0.6 s (m.a.d., 34 = experiments in 5 mice, **Figure 1**K). Overall, the time course of the estimated filter closely resembled the fast hemodynamic response function (HRF) measured optically in the cortex of awake mice (Pisauro *et al*., 2013); **Figure 1**L), with a possibly longer tail (Aydin *et al*., 2020). The estimated filter, therefore, likely corresponds to the hemodynamic response function (HRF) of the awake mouse cortex.

### Hemodynamic coupling across stimulus conditions and neural sources

This simple linear relationship explained cortical fUSI signals also during visually driven activity. To evoke visual responses, we presented a sequence of flashing checkerboards on the left, center, and right of the visual field (**Figure 2**A). In this sequence, there was only a 2.5% chance that a stimulus would reappear consecutively in the same position. The typical interval between stimuli in the same position was >7 s and often much longer, allowing fUSI signals to return to baseline between stimuli. An event-related analysis revealed the expected representation of visual space in both primary visual cortex and superior colliculus, with lateral stimuli driving fUSI responses in the opposite hemisphere, and the center stimulus driving both hemispheres (Brunner *et al*., 2020; Gesnik *et al*., 2017; Macé *et al*., 2018) (**Figure 2**B). Just as with spontaneous activity, the fUSI signal was well predicted by smoothing the firing rate with the estimated HRF (**Figure 2**C,D). Across experiments the median correlation between filtered firing rate and fUSI signals, in held-out data, was ρ = 0.55 ± 0.22 (m.a.d. across 34 experiments in 5 mice).

**Figure 2.**
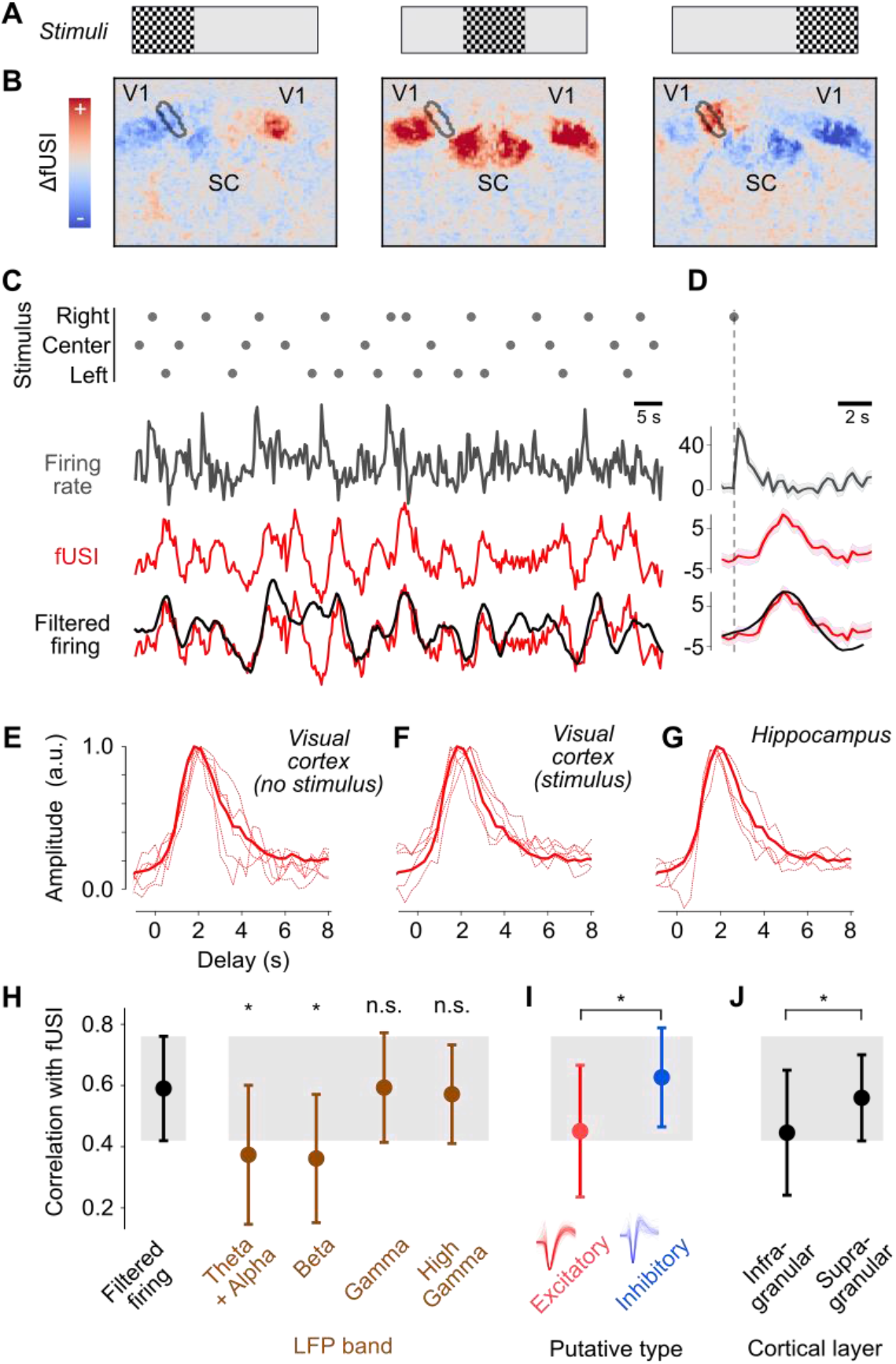
Hemodynamic coupling across stimulus conditions and neural sources. (**A**) Flashing checkerboards were presented on the left, center or right. (**B**) fUSI voxel responses to checkerboards, showing deviations from the mean activity. Black outline indicates the voxels traversed by the Neuropixels probe in V1. (**C**) Response to an example sequence of 30 stimuli (*dots*), showing firing rate in left V1 (*gray*), the corresponding fUSI signal (*red*), and the filtered firing rate (*black*). (**D**) Same format as C, showing the average response to the right (optimal) stimulus. (**E-F**). The estimated HRFs for visual cortex under spontaneous activity and visual stimulation for individual mice (n=5) resembled the mean HRF computed across mice, areas, and conditions (thick curve). (**G**). Individual HRFs for hippocampus estimated across spontaneous activity and visual stimulation (n=4) resembled the mean HRF (thick curve, same as in E-F). (**H**) Correlation between fUSI signals and LFP bands (n=187 recordings across hippocampus and visual cortex, in 5 animals). Asterisks indicate significant differences between firing rate and LFP bands (P<10^-12^). (**I**) Correlation between fUSI signals and putative excitatory and inhibitory neurons (n = 187 recordings). (**J**) Correlation between fUSI signals and infragranular and supragranular units recorded from visual cortex (n = 100 recordings in 5 animals).

The estimated HRF relating firing rate to fUSI signals was similar across mice and stimulus conditions, and between cortex and hippocampus. The HRFs measured in visual cortex in different mice were similar, both during spontaneous activity and during visual stimulation (**Figure 2**E,F). Moreover, they resembled the HRFs measured in hippocampus (**Figure 2**G). To assess whether the same HRF applies across mice, stimulus conditions (visual stimulation vs. spontaneous activity), and brain regions (visual cortex vs. hippocampus), we compared the predictions of fUSI signals obtained while allowing different HRFs vs. imposing a single average HRF (thick curve in **Figure 2**E-G). We used cross-validation to avoid over-fitting, and found that this single HRF explained a similar fraction of the variance as the individual HRFs. Therefore, though visual cortex and hippocampus show marked differences in vascular networks (Shaw et al., 2021), they have similar hemodynamic responses.

fUSI signals correlated equally well with neuronal firing rates and with local field potentials (LFP) in the gamma range. The LFP reflects average neural activity in a local region (Buzsáki *et al*., 2012; Katzner *et al*., 2009). We measured its power in four frequency bands: 4-12 Hz (alpha and theta), 12-30 Hz (beta), 30-90 Hz (gamma), and 110-170 Hz (high gamma). Consistent with previous findings (Aydin *et al*., 2020; Lima *et al*., 2014), fUSI signals correlated best with LFP signals in the gamma and high-gamma range (**Figure 2**H). These correlations were not significantly different from those observed with firing rates (P < 10^-12^, paired t-test). Correlations with the two lower LFP frequency bands, instead, were significantly lower (P = 0.57 and P = 0.08).

fUSI signals were best correlated with the activity of putative inhibitory neurons. We distinguished putative excitatory and inhibitory neurons based on their spike shape (Barthó et al., 2004), and filtered their firing rates separately with the estimated HRF. fUSI signals correlated significantly better with the filtered firing of putative inhibitory neurons than of putative excitatory neurons (ρ = 0.63 vs 0.45, P < 10^-12^, paired t-test). This difference was not due to a larger number of spikes, because the putative inhibitory neurons collectively fired fewer spikes. Indeed, the difference was still significant when we equated spike numbers by subsampling (P < 10^-12^).

In cortex, finally, fUSI signals were best correlated with activity measured in supragranular layers. This activity was significantly more correlated with fUSI signals than activity in infragranular layers (ρ = 0.56 vs 0.44, P = 0.005, paired t-test). Again, this effect was not due to larger numbers of spikes, because supragranular neurons have lower firing rates (Sakata and Harris, 2009). Indeed, the difference was still significant when we equated spike numbers through subsampling (P = 0.002).

### fUSI signals and firing rate are correlated across hemispheres

Consistent with previous results, fUSI signals showed broad spatial correlations: activity in one location was highly correlated with activity at nearby locations and in the opposite hemisphere. Similar to BOLD fMRI signals (Desjardins et al., 2001; Fox *et al*., 2007; Fox *et al*., 2006; Macey et al., 2004; Murphy et al., 2009), fUSI signals have broad spatial correlations within and across hemispheres, allowing the use of fUSI to study “functional connectivity” (Ferrier *et al*., 2020; Osmanski *et al*., 2014; Rabut *et al*., 2020; Rahal *et al*., 2020; Urban *et al*., 2015). Indeed, the fUSI signals measured in left visual cortex during spontaneous activity correlated highly with signals in many other cortical and subcortical locations, including those in the opposite hemisphere (**Figure 3**A,C). Correlations between fUSI signals across hemispheres were as high as ρ = 0.75 ± 0.08 (median ± m.a.d. across 68 recordings; **Figure 3**E, *left*). Similar results were seen in the hippocampus (**Figure 3**F,H), where the bilateral correlations were as high as ρ = 0.90 ± 0.04 (across 58 recordings; **Figure 3**J, *left*).

**Figure 3.**
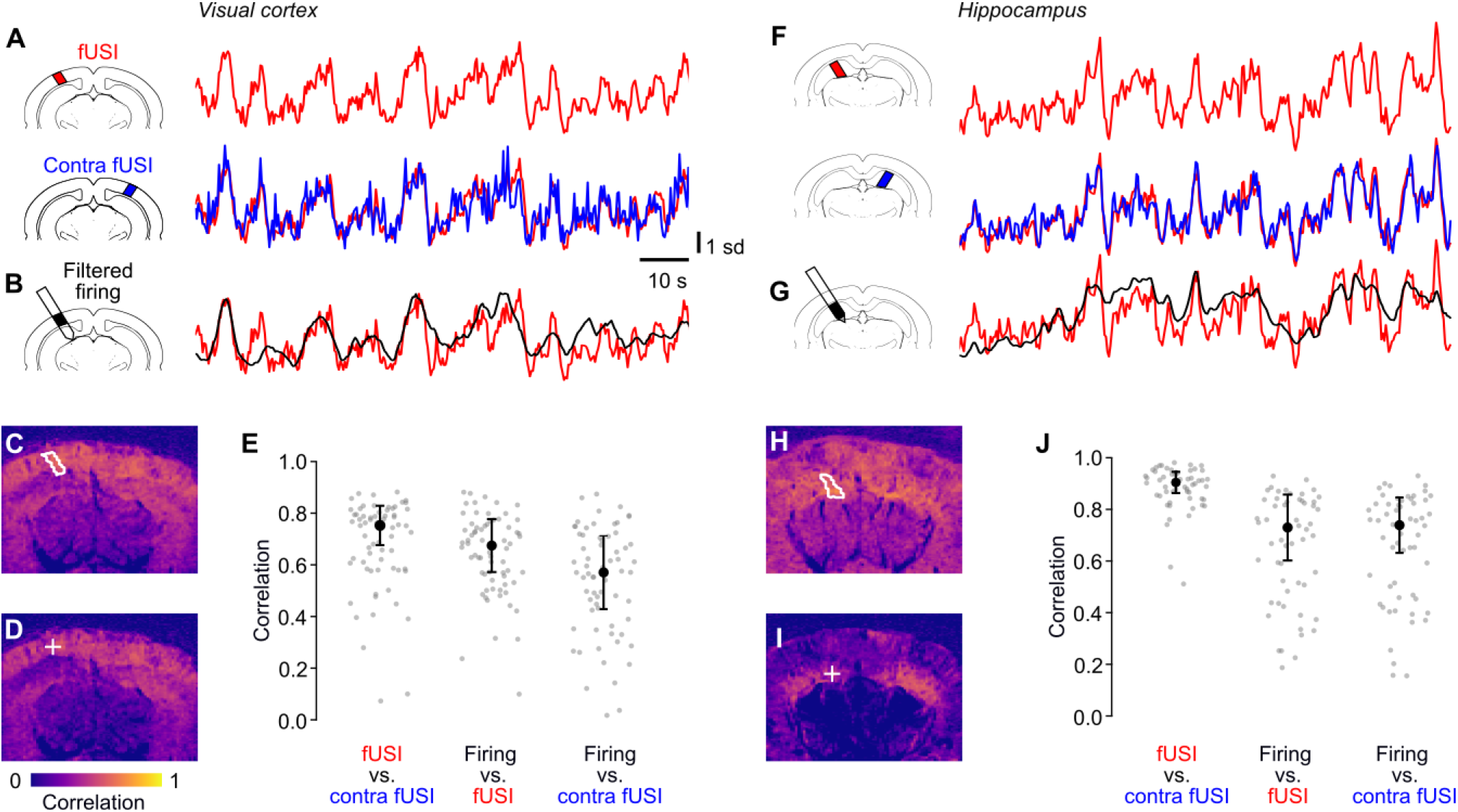
fUSI signals and firing rate are correlated across hemispheres. (**A**) fUSI traces measured during spontaneous activity in an example recording, in a ROI in the left visual cortex (*top*) and in a symmetrical ROI in right visual cortex (*bottom*). (**B**). Filtered firing rate measured simultaneously in the left ROI. (**C**) Correlation between the fUSI voxels in the left ROI (*white contour*) and all the individual fUSI voxels. (**D**) Correlation between the filtered firing rate measured in the left ROI (*plus sign*) and all the individual fUSI voxels. (**E**) Correlations between fUSI signals in left and right visual cortex (*left*) and between filtered firing rate and simultaneous fUSI signals in the same location in visual cortex (*center*) and in the opposite hemisphere (*right*). Black dot and error bars show median ± m.a.d across n = 68 recordings during spontaneous activity and visual stimulation. (**F-J**) Same analyses for recordings where firing rate and fUSI were measured in hippocampus (n = 58 recordings).

Accordingly, the filtered firing rate correlated not only with fUSI signals at the same location but also at other locations, including those in the opposite hemisphere. The filtered firing rate measured in left visual cortex resembled fUSI signals measured in the same location and in the opposite hemisphere (**Figure 3**B,D). Correlations with contralateral fUSI signals were ρ = 0.57 ± 0.14, barely lower than correlations with ipsilateral fUSI signals (ρ = 0.68 ± 0.10, **Figure 3**E, *center* and *right*). Likewise, the filtered firing rate measured in left hippocampus resembled fUSI signals measured in both left and right hippocampus (**Figure 3**G,I), with correlations above 0.7 in both cases (**Figure 3**J).

These results suggest that the strong spatial correlations seen in fUSI signals may be explained by underlying bilateral fluctuations in neural activity. Indeed, ongoing neural activity has broad spatial correlations and is strongly bilateral, both during rest and during behavior (Mohajerani *et al*., 2010; Musall *et al*., 2019; Shimaoka *et al*., 2019). However, there is another possible source of broad, bilateral correlations: perhaps there are hemodynamic fluctuations that are broad and bilateral but unrelated to neuronal activity (Drew et al., 2020; Turner *et al*., 2020).

### Bilateral firing largely explains bilateral fUSI signals

To investigate the high bilateral correlations observed in fUSI we performed simultaneous recordings with two Neuropixels probes. In three of the mice, we inserted two probes symmetrically relative to the midline, targeting bilateral locations in visual cortex (**Figure 4**A). We could thus not only compare filtered firing rates to fUSI signals (**Figure 4**A) and fUSI signals across hemispheres (**Figure 4**B), but also firing rates measured across hemispheres (**Figure 4**C).

**Figure 4.**
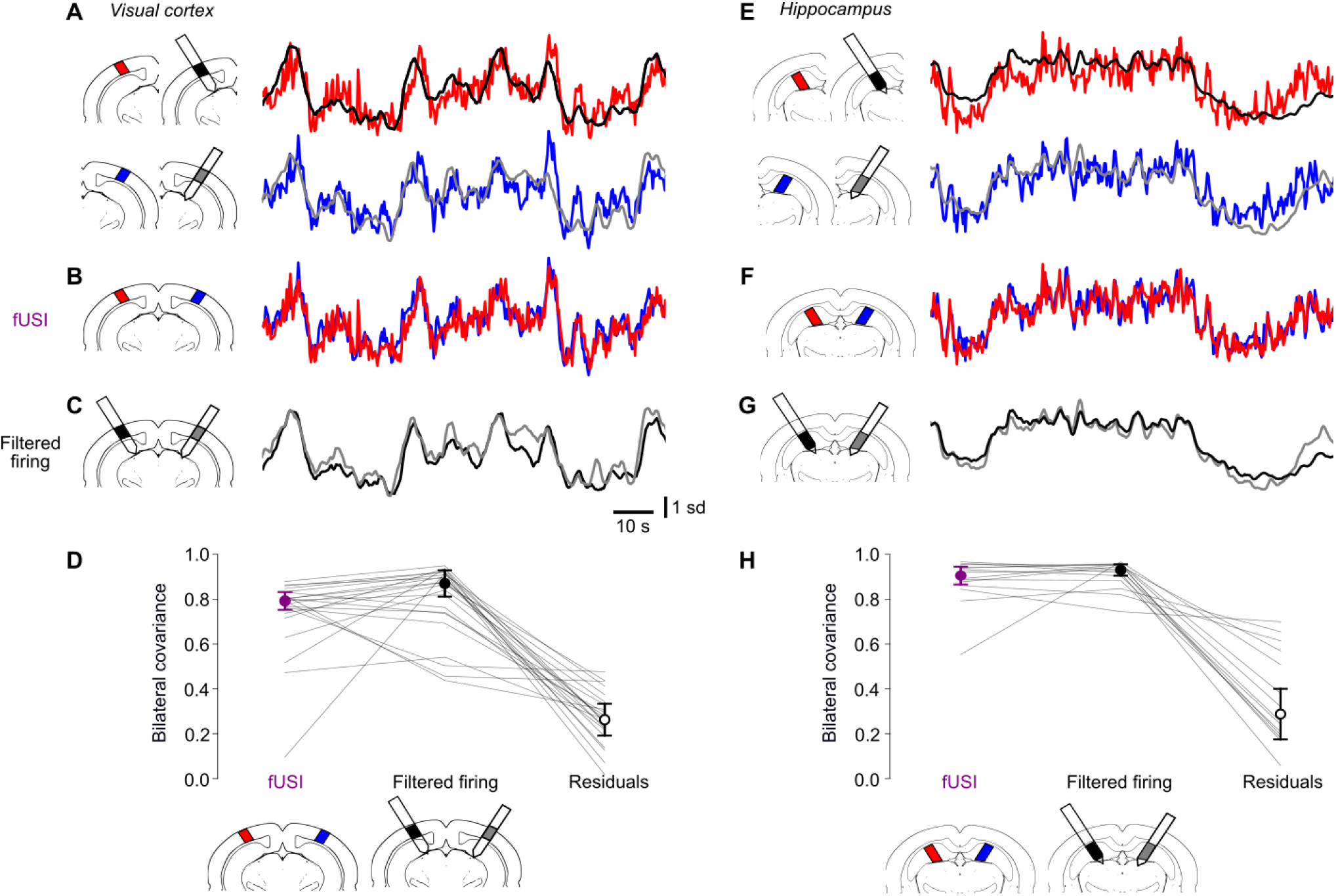
Bilateral firing largely explains bilateral fUSI signals. (**A**) Example recordings from two Neuropixels probes inserted bilaterally, yielding simultaneous measurements of firing rate (filtered with the HRF, *black* and *gray* curves) and fUSI signals (*red* and *blue* curves) during spontaneous activity in left and right visual cortex. (**B**,**C**) Superposition of the bilateral fUSI signals (**B**) and of the bilateral filtered firing rates (**C**). (**D**) Covariance between left and right fUSI signals (*left*), filtered firing rates (*middle*), and residuals obtained by subtracting the filtered firing rates from the fUSI signals (*right*). Because fUSI signals and filtered firing rates are z-scored, for these two quantities, covariance is equal to correlation. *Dots* and *error bars* indicate median ± m.a.d. across 22 recordings (*lines*) in 3 mice during spontaneous activity and visual stimulation. (**E-H**). Same analysis, for hippocampus (14 recordings in 3 mice).

The filtered firing rates in left visual cortex closely resembled those simultaneously recorded in right visual cortex (**Figure 4**C). Across recordings, the two filtered firing rates had high bilateral correlation, ρ = 0.87 ± 0.06 (median ± m.a.d., across 22 recordings; **Figure 4**D, *middle*). These correlations in firing rate matched those measured in fUSI signals (**Figure 4**D, *left*), which were not significantly higher (paired t-test P = 0.28, n = 22).

Similar results were seen in the left and right hippocampus (**Figure 4**E-G). The filtered firing rate measured in left and right hippocampus exhibited strong bilateral correlation, ρ = 0.93 ± 0.03 (median ± m.a.d. across 14 recordings; **Figure 4**H, *middle*). These bilateral correlations were not significantly lower than those measured in fUSI signals (**Figure 4**H, *left*, P = 0.40, n = 14).

To test whether the bilateral correlations in firing rates fully explain the bilateral correlations observed in fUSI, we removed the fluctuations in fUSI signals that were predicted by the filtered firing rate measured at the same location and examined the residuals. The residuals had much smaller bilateral covariance than the original fUSI signals, both in visual cortex (paired t-test P < 10^-10^, **Figure 4**D, *right*), and in hippocampus (P < 10^-4^, **Figure 4**H, *right*). These fUSI residuals strongly correlated across the entire fUSI slice (**Suppl. Figure 2**), suggesting that they reflect micromovements of the brain and global vascular effects. The latter may include aliasing of small respiratory and heartbeat movements, and spontaneous oscillations in arterial diameter (Drew, 2019; Winder *et al*., 2017).

## Discussion

Much of brain activity is endogenous – unrelated to external events – so it must be measured in individual trials. Single-trial measurements of brain activity, indeed, are routine with electrophysiology techniques that record local neuronal spikes. However, they are exceedingly difficult with methods that have larger spatial coverage, such as fMRI and EEG. These methods have low signal/noise ratios, and thus require recordings to be averaged across trials (event-related analysis) or internal events (e.g., correlation with a seed voxel).

Our results indicate that functional ultrasound imaging (fUSI) in mice can bridge this gap, providing large-scale measurements of brain activity in single trials. By performing simultaneous electrophysiology and fUSI, we were able to establish the relationship between neuronal firing and ultrasound signals on a trial-by-trial, moment-by-moment basis. The results indicate that functional ultrasound signals measured at frequencies below 0.3 Hz strongly correlate with neural activity. Indeed, thanks to their high signal/noise ratio, fUSI signals can even drive brain-machine interfaces (Norman et al., 2021).

We found that fUSI signals bear a simple relationship to the underlying neural activity captured by convolution with a standard hemodynamic response function. These results confirm and extend previous work that related blood signals to fUSI measurements performed separately and averaged across trials (Aydin *et al*., 2020; Boido *et al*., 2019). They suggest that the hemodynamic response function measured with fUSI is the same that had been measured optically (Pisauro *et al*., 2013), and is consistent across mice, stimulus conditions, and brain regions. However, we only tested two brain regions – visual cortex and hippocampus – and further investigations might reveal different hemodynamic responses elsewhere in the brain (Handwerker et al., 2004).

fUSI signals may appear noisy because they are variable in time and broadly correlated in space. However, this reflects not just measurement error, but true structured fluctuations in neuronal firing. Brain activity involves the simultaneous firing of large numbers of neurons across regions. These broad activations are typically associated with fluctuations in internal state and body movement (Drew *et al*., 2019; Musall *et al*., 2019; Stringer *et al*., 2019), and can be highly correlated across hemispheres (Drew *et al*., 2019; Fox *et al*., 2007; Fox *et al*., 2006; Mohajerani *et al*., 2010; Shimaoka *et al*., 2019). Accordingly, our double recording experiments reveal that fUSI signals match neural activity even when they spread over large portions of the brain, including the opposite hemisphere.

We found a correlation as high as 0.6 between fUSI signals and smoothed firing rates in mice that were mainly awake and alert. The correlation might be even higher if it were measured during NREM sleep, when the relationship between blood flow and neural activity is thought to be particularly strong (Turner et al., 2020).

fUSI signals correlated best with the firing of fast-spiking, putative inhibitory neurons. This observation may relate to a specific role of synaptic inhibition in controlling blood flow (Anenberg et al., 2015; Cauli et al., 2004). However, fast-spiking cells are likely to be largely parvalbumin-positive interneurons, whose activation reduces, rather than increase, blood flow (Lee et al., 2021). The high correlation with inhibitory activity seems thus more likely to arise because inhibitory neurons are robust estimators of nearby firing rate (Isaacson and Scanziani, 2011), pooling over more neurons than those surrounding the probe.

Perhaps a similar reasoning explains the higher correlations we observed between fUSI signals and activity in supragranular layers of the cortex. These laminar differences were small but significant, and may make it difficult to measure laminar activity with fMRI (Huber et al., 2017)

By releasing the data from our simultaneous recordings and fUSI imaging (*URL to go here upon publication*), we hope to facilitate improvements to the fUSI processing pipeline. This pipeline begins from raw Doppler images and aims to isolate signals related to neural activity from noise originating, e.g., from tissue movement (Baranger *et al*., 2018; Demené *et al*., 2015; Macé *et al*., 2011; Macé *et al*., 2013). It involves multiple steps, including temporal high-pass filtering, principal component analysis, and subsequent removing of the largest components. We confirmed that the present pipeline is adequate: it yields fUSI signals that are closely related to the underlying firing rates. However, it may be amenable to further improvements. Moreover, it should be possible to design a deconvolution filter that estimates firing rate from fUSI signals, much as one can estimate firing rates from widefield calcium fluorescence (Peters et al., 2021). To validate all this, it is essential to have neuronal spikes as ground-truth data.

We conclude that fUSI signals bear a simple relationship to neuronal firing and accurately reflect this firing both in time and in space. We hope that these results will be useful to the increasing numbers of laboratories that use functional ultrasound imaging to reveal brain function.

## Acknowledgments

We thank Wang Qi for help with brain alignment and pupil detection, and Célian Bimbard for helpful comments. This work was funded by a joint Wellcome Trust Investigator Award to MC and KHD (grant 205093/Z/16/Z). AU and GB were supported by grants from the Leducq Foundation (15CVD02), FWO (MEDI-RESCU2-AKUL/17/049, G091719N, and 1197818N), VIB Tech-Watch (fUSI-MICE), and the internal techfund from Neuro-Electronics Research Flanders (NERF). MC holds the GlaxoSmithKline / Fight for Sight Chair in Visual Neuroscience.

## Author Contributions

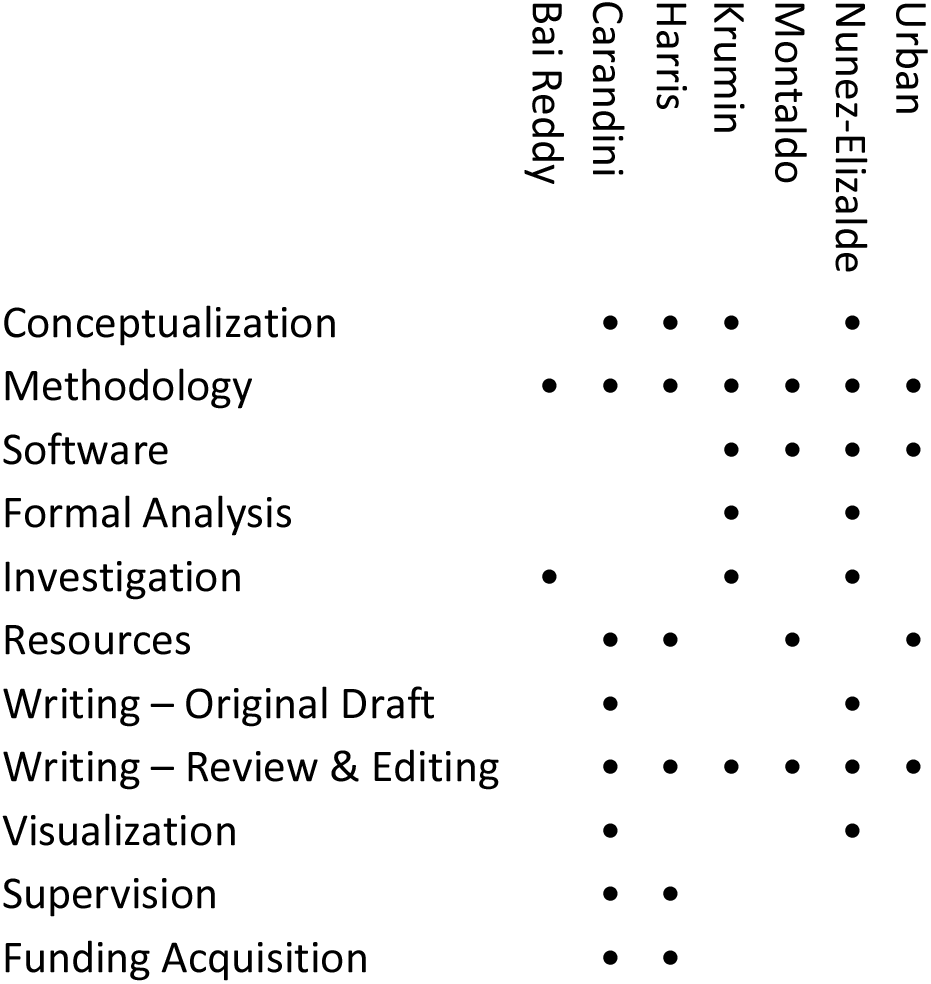

## Methods

All experimental procedures were conducted according to the UK Animals Scientific Procedures Act (1986). Experiments were performed at University College London, under a Project License released by the Home Office following appropriate ethics review.

### Initial surgery

Experiments were conducted in 5 C57/BL6 mice (4 male, 1 female), 9-12 weeks of age. Mice were first implanted with a headplate and cranial window under surgical anesthesia in sterile conditions. Procedures for implanting the headplate are standard in the field (e.g., International Brain Laboratory et al., 2021). The cranial window replaced a dorsal section of the skull (~8 mm in ML and ~5 mm in AP) with 90 μm thick ultrasound-permeable polymethylpentene (PMP) film (ME311070, Goodfellow Cambridge Ltd.). The PMP film was then covered with Kwik-Cast (World Precision Instruments, USA), except during imaging sessions. This initial surgery was followed by 5-12 days of recovery, handling, and habituation to the experimental rig.

### Recording sessions

In each recording session, we head-fixed the mice by securing the headplate to a post placed 10 cm from three computer screens (Adafruit, LP097QX1, 60Hz refresh rate) arranged at right angles to span 270 deg in azimuth and ~70 deg in elevation. Fresnel lenses (f = 220 mm, BHPA220-2-5, Wuxi Bohai Optics) were mounted in front of the screens to reduce intensity differences across parts of the screens that are viewed from different angles. The lenses were covered with diffusing film (Frostbite, The Window Film Company) to reduce specular reflections.

We then inserted a Neuropixels probe (Jun *et al*., 2017) through a hole in the PMP film (0.5 mm radius). The probe described a parasagittal trajectory (posterolateral to anteromedial), at an angle of 28 deg relative to the midline (sagittal plane) and 40 deg relative to the horizontal (axial) plane. In some experiments we introduced a second Neuropixels probe in the opposite hemisphere, along the mirror-symmetric trajectory.

We then covered the PMP film with ultrasound gel and positioned an ultrasound transducer above it (128-element linear array, 100 μm pitch, 8 mm focal length, 15 MHz central frequency, model L22-Xtech, Vermon, France). Doppler signals from the transducer were acquired using a Vantage 128 ultrasound system (Verasonics, USA) controlled by a custom Matlab-based user interface (Alan Urban Consulting) recording continuously at 500 Hz. fUSI acquisition was synchronized with the visual stimulus by recording the TTL pulses of the fUSI frames together with the flickering sync square on the visual stimulus monitor (using TimeLine, Bhagat et al., 2020). A similar method was used to align the Neuropixels recordings, by simultaneous recording external TTL pulses on an additional channel of a Neuropixels probe and on TimeLine.

In each recording session, we moved the ultrasound transducer to cover 3-5 coronal slices. For each slice, we performed two recordings: first, we displayed a gray screen for ~4 minutes to measure spontaneous activity; second, we presented flashing checkerboards flashing at 2 Hz for ~8 minutes to measure stimulus-evoked responses. The checkerboards were presented in the left, center, or right screens (one screen at a time). Checkerboard squares had a size of 15 deg and could be white or black. The checkerboard sequence was interspersed with blank trials. The sequence consisted of 40 checkerboards, lasted ~90 seconds and was repeated 4-5 times.

At the end of the recording session, we slowly extracted the Neuropixels probe from the brain while recording fUSI images from one coronal slice. This movement allowed us to localize the probe’s tip within the fUSI slice, giving us a 2D coronal projection of the probe’s 3D trajectory.

Finally, we acquired a series of coronal fUSI images (a “Y-stack”) from posterior to anterior, spaced 0.1 mm apart. These images were later used to construct a 3D fUSI volume of the brain to facilitate registration with the Allen Atlas and to identify the location of the Neuropixels probe in the fUSI slices.

### Processing of ultrasound signals

fUSI signals were computed using standard methods (Macé *et al*., 2011). The 500 Hz complex-valued Doppler signals were divided into 400 ms chunks that overlapped by 50 ms. Then, each chunk was high-pass filtered with a cut-off of 15 Hz, and its principal components were computed in space and time. The first 15 principal components were then removed, to remove artifacts including those due to brain movement (Demené *et al*., 2015). A power Doppler image was then computed by squaring the complex-valued signals and averaging them in the central (non-overlapping) 300 ms window, for a final temporal resolution of 3.33 Hz. The voxel time courses were then converted to fractional change relative to the mean of each voxel.

We computed the fUSI signal trace for a region of interest (ROI) by taking the mean of the individual time courses of voxels in the ROI. The individual voxel time courses were normalized to percent signal change units before computing their mean.

fUSI images were manually aligned to a vascular atlas with Allen CCF labels (Todorov *et al*., 2020). We first registered the 3D volume from each recording session to the vasculature atlas. To this end, we used FreeSurfer (Fischl, 2012) to rotate, shift, and scale the vasculature atlas to match the vasculature features salient in the fUSI 3D volume. Once aligned, the transformation relating the vasculature atlas to the fUSI volume was saved and applied to the vasculature-matched Allen CCF labels. Finally, the Allen CCF labels were resampled to match the spatial resolution of the fUSI volume (100 x 100 x 48 μm^3^), yielding Allen CCF labels for each fUSI voxel.

### Spatial alignment

To identify brain locations simultaneously traversed by the Neuropixels probe and the fUSI slices, we estimated the 3D trajectory of the Neuropixels probe in the fUSI Y-stack volume. Based on the geometry of the simultaneous recordings, we located the Neuropixels probe insertion site ~0.2 mm behind the posterior-most fUSI slice. We then reconstructed the Neuropixels probe 3D trajectory so that its 2D coronal projection best matched the 2D coronal projection measured with fUSI *in vivo* during Neuropixels probe extraction. This 3D trajectory allowed us to map from Neuropixels probe sites to fUSI voxels in a slice, and vice versa.

While the Neuropixels probe intersects with the fUSI slice plane at one point, the fUSI slice has a thickness. This thickness has a full-width at half maximum of ~300 μm (Brunner *et al*., 2020) and not larger than 500 μm (Demené et al., 2016). The fUSI voxels and Neuropixels probe sites located 250 μm on either side of the fUSI plane (along the Y-axis) were used for the analyses.

For each recording, we identified the fUSI voxels that were intersected by the Neuropixels probe and used them to define a region of interest (ROI). ROIs for visual cortex and for hippocampus tended to have a similar number of voxels (~50 voxels). The fUSI signal within the ROI was computed as the mean of the individual voxel time courses.

### Processing of electrophysiological signals

The electrophysiology data was spike sorted using kilosort2 (Pachitariu et al., 2016) and the resulting output was then manually curated with Phy (github.com/cortex-lab/phy). Manual curation sought to identify clusters corresponding to single- and multi-unit activity and to remove spurious and noisy clusters based on traditional measures such as inter-spike interval, autocorrelation, waveform shape. After spike sorting, single- and multi-unit activity was summed across the electrode sites that traversed the fUSI imaging plane to obtain a single firing rate trace for the ROI. This trace was binned at 300 ms intervals to match the temporal resolution of fUSI signals. To distinguish spikes of putative excitatory and inhibitory neurons we clustered based on spike width (Barthó *et al*., 2004; Lin et al., 2020).

To analyze the LFP signals we took the LFP output of the Neuropixels probes and separated it into four frequency bands using established methods (Lima *et al*., 2014).

To identify the Neuropixels probe sites located in visual cortex and in hippocampus, we used the cross-correlation of the multi-unit activity. We divided the Neuropixels probe sites into non-overlapping 100 μm segments and computed their cross-correlation. Sites at the top of the Neuropixels probe corresponded to visual cortex and were strongly correlated with each other. Sites immediately below the visual cortex corresponded to the hippocampus and were strongly correlated with each other.

To obtain ROIs in the fUSI images we identified the fUSI voxels traversed by the Neuropixels probe in visual cortex and hippocampus using the probe’s 3D trajectory and a labeled volume of the standard C57 mouse brain, the Allen Common Coordinate Framework (CCF, Wang *et al*., 2020). For a ROI in visual cortex or hippocampus we included all voxels that were (1) in the fUSI slice; (2) in the appropriate brain region according to the CCF; and (3) in the Neuropixels probe trajectory.

### Cross-correlation and coherence

The cross-correlation between firing rate and fUSI signal traces was computed at different delays by shifting the firing rate relative to the fUSI signals (from –5 to +30 s).

Coherence was computed using the multi-taper method (github.com/nipy/nitime). To do this, we used three minutes of firing rate and fUSI signal traces recorded simultaneously during periods of spontaneous activity. We computed the coherence between signals up to 1.667 Hz, the Nyquist limit of our 300 ms sampling interval.

To compute the chance coherence between fUSI signals and firing rate, we randomly and circularly shifted the firing rate and computed its coherence with the original fUSI signal trace. This process was repeated 1,000 times and computing the mean at each frequency. The chance coherence was then computed as the median across recordings for each frequency.

To determine the highest frequency at which firing rate and fUSI signals are coherent, we compared the actual versus chance coherence values across sessions. To do this, we found the frequencies at which coherence was above chance. We then identified the highest of these frequencies (0.32 Hz). Above this frequency, the coherence between firing rate and fUSI matches what can be expected by chance.

### Hemodynamic response function

To estimate the hemodynamic response function relating firing rate to fUSI, we modelled fUSI responses for each recording as a convolution of the firing rate in time with a finite impulse response filter. The optimal filter for each recording was estimated using cross-validated ridge regression (Hoerl and Kennard, 1970) using open-source software (Nunez-Elizalde et al., 2019). To avoid overfitting, the data were split into a training and a test set (75%/25%). Using the training set, the optimal regularization parameter was found independently in each recording using a 5-fold cross-validation procedure twice. The accuracy of the model was assessed by computing the correlation between predicted and actual fUSI signals in the held-out test set. Finally, the hemodynamic response function was estimated for each recording using 100% of the data.

### Whisker movements and pupil diameter

To assess alertness, we recorded videos of the mouse’s face during our experiments (**Suppl. Figure 2**). Using these videos, we quantified pupil size and whisker motion. Pupil diameter was estimated with DeepLabCut (Mathis et al., 2018; Meijer et al., 2020). Whisker motion was estimated following established procedures (Stringer *et al*., 2019) using a pyramid of spatiotemporal Gabor filters (github.com/gallantlab/pymoten). The difference between frames was computed for each pixel, yielding a time-by-pixels matrix. The principal components were then computed by concatenating all frames, and the top 10 components were used to compute the total energy over time.

## Supplementary Figures

**Supplemental Figure 1.**
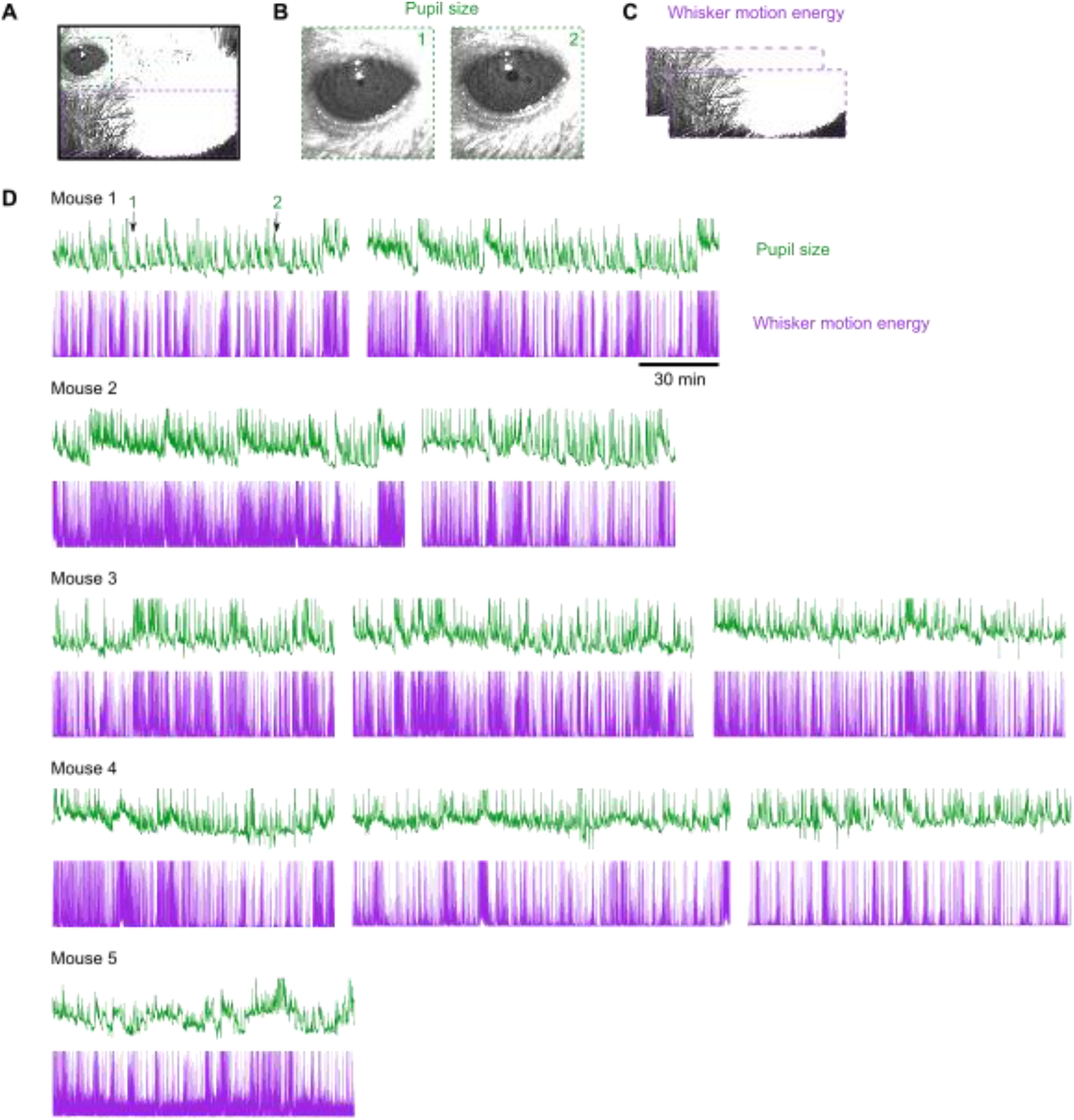
Behavioral monitoring. (**A**) Example frame from a camera pointed at the mouse face, showing regions analyzed for eye (*green*) and whiskers (*purple*). (**B**) Example frames of the eye, used to estimate pupil size, showing a frame with smaller pupil (*1*) and one with larger pupil (*2*). (**C**) Example frames of the whiskers, used to estimate whisker motion energy. (**D**) Pupil size (*green*) and whisker motion energy (*purple*) for 11 recording sessions in 5 mice. Arrows 1 and 2 mark the frames in B.

**Supplemental Figure 2.**
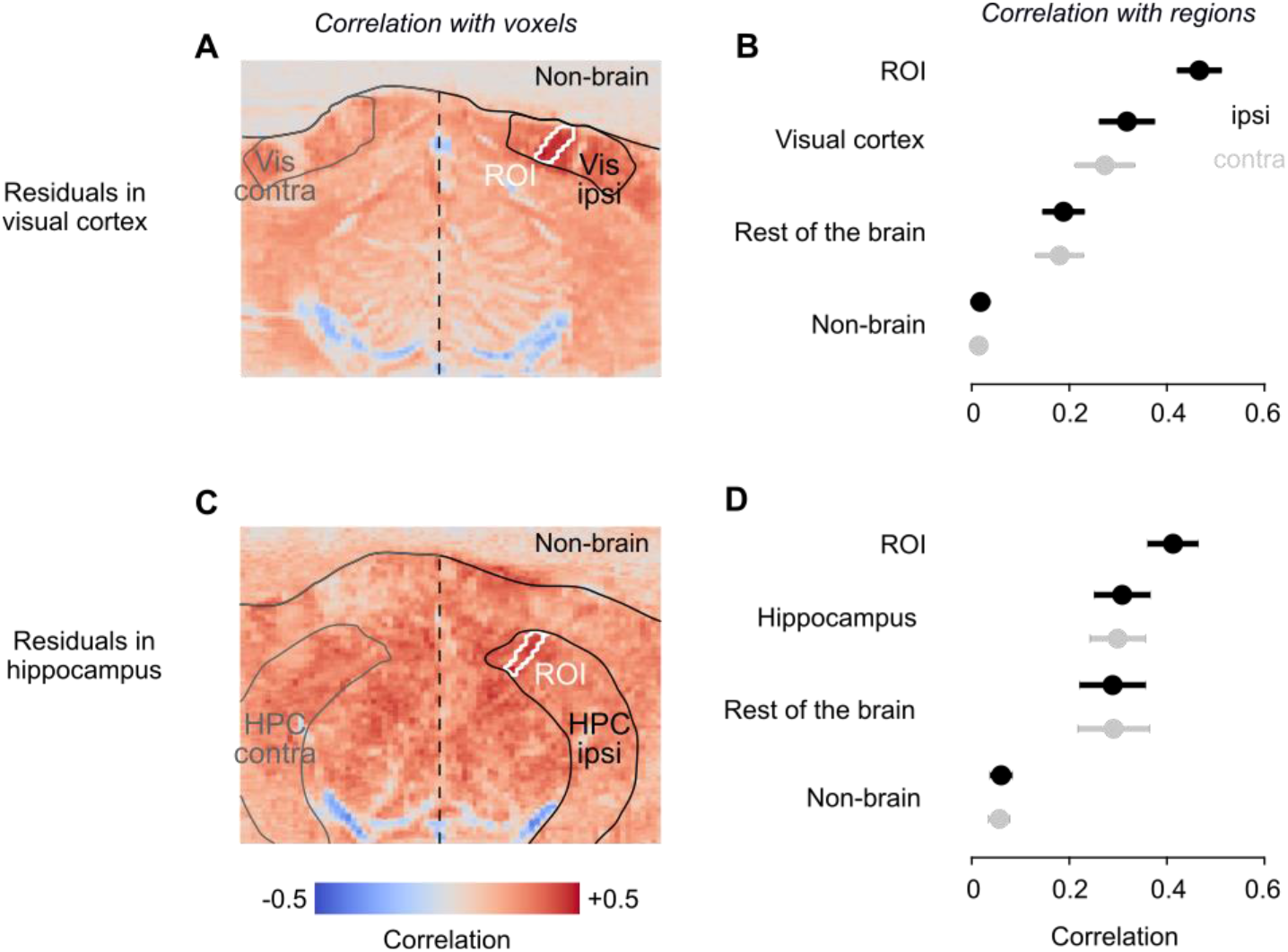
fUSI residuals correlate with the whole brain. (**A**) Correlations between fUSI residuals (fUSI signals minus filtered firing rate) in visual cortex with fUSI signals in the whole slice. (**B**) Correlations with fUSI signals in the ROI, in the rest of visual cortex, in contralateral visual cortex, and in the rest of the brain. (**C**,**D**) Same, for fUSI residuals in hippocampus.

